# Diurnal rhythms in durum wheat triggered by *Rhopalosiphum padi* (bird cherry-oat aphid)

**DOI:** 10.1101/2024.08.25.609566

**Authors:** Yoshiahu Goldstein, Jinlong Han, Daniel Kunk, Albert Batushansky, Vamsi Nalam, Vered Tzin

**Affiliations:** French Associates Institute for Agriculture and Biotechnology of Drylands, Jacob Blaustein Institutes for Desert Research, Ben-Gurion University of the Negev, Midreshet Ben Gurion, Israel; Department of Agricultural Biology, Colorado State University, Fort Collins, CO, USA; Department of Cell and Molecular Biology, Colorado State University, Fort Collins, CO, USA; Ilse Katz Institute for Nanoscale Science & Technology, Ben-Gurion University of the Negev, Beer Sheva, Israel

**Keywords:** Rhythmicity, diurnal, WRKY, transcription factor, transcriptomics, untargeted metabolomics, wheat, *Rhopalosiphum padi*, aphid

## Abstract

Wheat is a staple crop and one of the most widely consumed grains globally. Wheat yields can experience significant losses due to the damaging effects of herbivore infestation. However, little is known about the effect aphids have on the natural diurnal rhythms in plants. Our time-series transcriptomics and metabolomics study reveal intriguing molecular changes occurring in plant diurnal rhythmicity upon aphid infestation. Under control conditions, 15,366 out of the 66,559 genes in the tetraploid wheat cultivar Svevo, representing approximately 25% of the transcriptome, exhibited diurnal rhythmicity. Upon aphid infestation, 5,682 genes lost their rhythmicity, while additional 5,203 genes began to exhibit diurnal rhythmicity. The aphid-induced rhythmic genes were enriched in GO terms associated with plant defense, such as protein phosphorylation and cellular response to ABA and were enriched with motifs of the WRKY transcription factor families. Conversely, the genes that lost rhythmicity due to aphid infestation were enriched with motifs of the TCP and ERF transcription factor families. While the core circadian clock genes maintain their rhythmicity during infestation, we observed that approximately 60% of rhythmic genes experience disruptions in their rhythms during aphid infestation. These changes can influence both the plant’s growth and development processes as well as defense responses. Furthermore, analysis of rhythmic metabolite composition revealed that several monoterpenoids gained rhythmic activity under infestation, while saccharides retained their rhythmic patterns. Our findings highlight the ability of insect infestation to disrupt the natural diurnal cycles in plants, expanding our knowledge of the complex interactions between plants and insects.

## Introduction

Plants, as sessile organisms, must activate an arsenal of defenses to cope with environmental challenges, including both abiotic and biotic stressors. They must continuously adapt to changing environmental conditions, including the diurnal cycle, which consists of alternating light (day) and dark (night) periods. This diurnal cycle is regulated by both external stimuli and by the plant’s internal clock, its circadian clock (Nozue and Maloof, 2006). These diurnal cycles in plants exhibit a 24-hour rhythmic pattern that is crucial for the plant’s growth and survival. Moreover, the plant circadian clock plays an important role in a plant’s defense system (McClung, 2006; Roden and Ingle, 2009).

Wheat is a staple crop consumed worldwide, contributing an estimated 20% of the total human protein intake (Erenstein et al., 2022). It maintains a circadian clock that influences growth by regulating major processes such as photosynthesis and several yield-related traits (Sun et al., 2020; Gong et al., 2022). In addition to playing a role in stress response defenses, its circadian clock also exhibits circadian gating in response to cold (Ren et al., 2023; Graham et al., 2023).

Insect herbivores also exhibit distinct daily feeding patterns, leading to an intriguing dynamic: plants synchronize their defenses with the diurnal activity of these herbivores. Goodspeed et al. (2012) provide evidence of this synchronization in the Arabidopsis-cabbage looper (*Trichoplusia ni*) system, focusing on jasmonate phytohormone-dependent defenses (Goodspeed et al., 2012). However, the broader implications of this synchronization, and the extent to which it influences the co-evolutionary arms race between plants and their herbivore antagonists, remain largely unexplored.

Aphids are phloem-sucking insects that cause major reductions in crop yield through metabolite consumption and through virus transmission (Kieckhefer and Gellner, 1992). With a wide variety of species, aphids can cause damage to a broad range of plants, including staple crops such as barley, maize, and wheat. Using their stylets, aphids pierce plant cells, causing minimal mechanical damage, then direct these stylets to the phloem cells for the ingestion of sap (Will and Van Bel, 2006). Aphids release salivary effectors, which are small, secreted molecules, into plant cells to modulate a plant’s physiological responses. Some effectors have been shown to inhibit plant defenses, thereby promoting the aphids’ success (Escudero-Martinez et al., 2021; Zhang et al., 2022). To better suit their nutritional needs, aphids can modify the phloem-sap composition for their benefit. For instance, they can increase the amount of free essential compounds, such as amino acids, in the predominantly sugar-rich phloem sap (Jakobs *et al*., 2019). *Rhopalosiphum padi* (bird cherry-oat aphid) is a major economic pest in wheat production and other important cereals (Voss *et al*., 1997). Beyond diminishing the plant’s metabolites when feeding, *R. padi* can also transmit the highly destructive barley yellow dwarf virus (Jiménez-Martínez and Bosque-Pérez, 2004). Like plants, aphids have their own diurnal cycle. *R. padi* shows increased time spent in the phloem during the night, possibly to regulate their osmotic potential (Nalam *et al*., 2021; Han *et al*., 2024). In addition, we recently showed that many putative aphid effectors exhibit diurnally rhythmic expression patterns, which could potentially affect the plant’s diurnal cycle. (Han *et al*., 2024).

Transcription factors are regulatory proteins that control gene expression by binding to promoter sites of specific DNA sequences and turning genes on or off. As transcription regulators, they play crucial roles in regulating both the plant circadian clock and plant responses to different stresses (Ng et al., 2018; Nakamichi, 2020). Some transcription factors have also been shown to interact with salivary effectors secreted into the plant. SmCSP4, an aphid salivary effector, directly interacts with the TaWRKY76 transcription factor, thereby activating a phytohormone degradation gene, leading to higher salicylic acid (SA) levels and lower aphid performance (Zhang *et al*., 2023). Considering their roles in regulating diurnal rhythms and their response to effectors, transcription factors might influence diurnal defense responses under aphid infestation.

In this study, we present a time-series analysis to understand the effect of aphid infestation on diurnal cycles in plants. Emphasis is placed on transcription factors due to their importance in regulating circadian rhythms and stress responses in plants. The gain and loss of gene expression rhythmicity due to aphid infestation, along with the associated biological pathways, are discussed.

## Results

### Rhythmic plant genes under control and aphid infestation

To evaluate the variability of the different timepoints, principal component analysis (PCA) was conducted on control (no aphids) and aphid-treated samples (Figures 1b and 1c, respectively). A clear separation is observed between the different time points of the two conditions, and for the control conditions, the first (12:00) and last (12:00) time points are in close proximity to each other.

**Figure 1.**
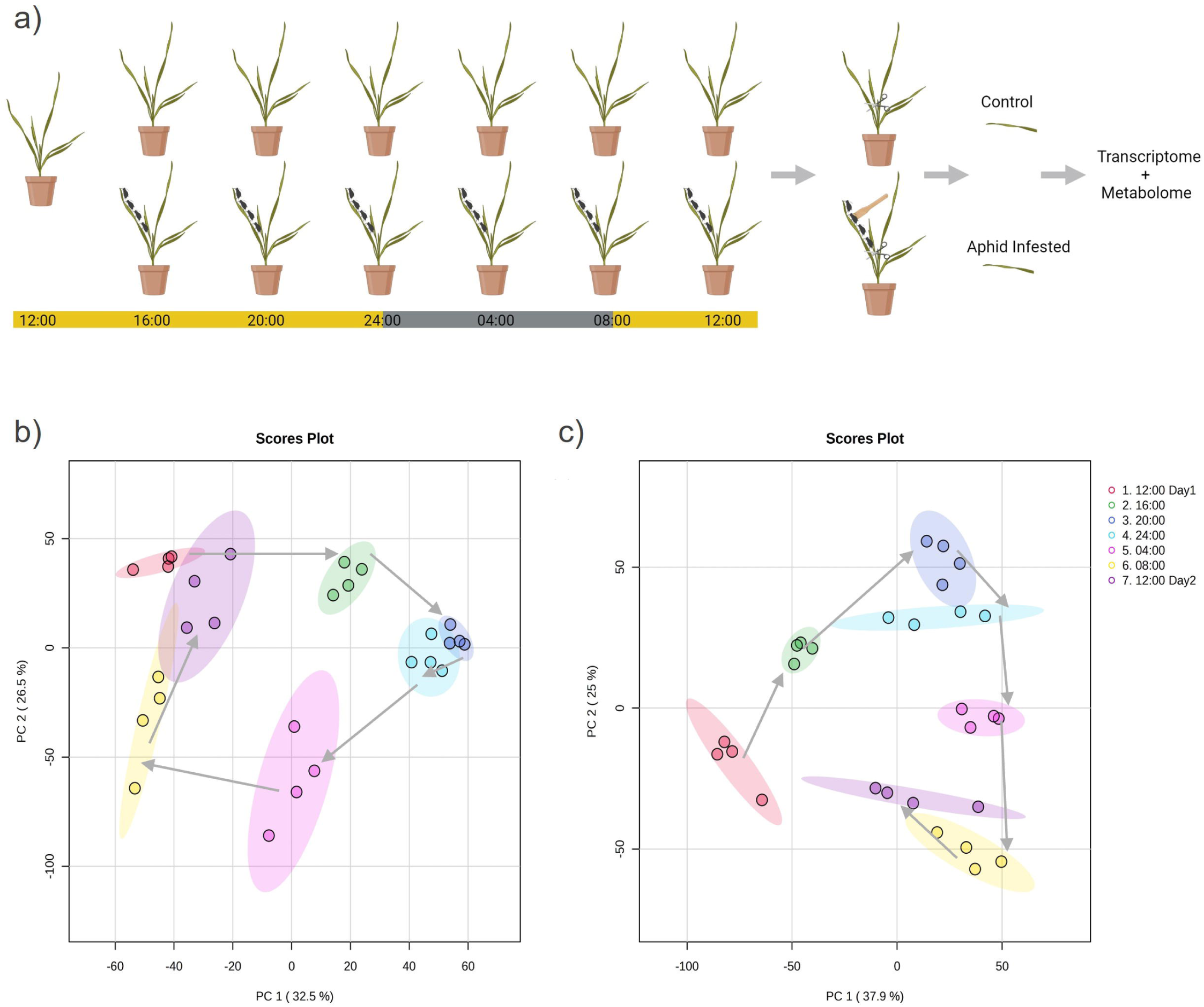
Experimental design and exploratory data analysis. (a) The experimental design was as follows: Svevo plants were infested with *R.padi* aphids at zt4 (12:00). Every 4 hours, samples were collected from both infested and control plants, with aphids gently swept off the leaves prior to collection. Samples were immediately frozen with liquid nitrogen and stored in −80°C for future processing. Four samples were collected for RNA-seq and six for untargeted metabolomics. (b) A PCA plot of control condition RNA-seq results was created using normalized expression levels with MetaboAnalyst, applying a 25% Interquartile Range (IQR) filter and Auto-scaling. Overall, PC1 has 32.5% and PC2 has 26.5% variance. Timepoints follow a clockwise position, with the first sampling collected at 12:00 on the first day in close proximity to the last sample collected at 12:00 on the second day. (c) A PCA plot of aphid-infested condition genes was created using normalized expression levels with MetaboAnalyst. Overall, PC1 has 37.0% and PC2 has 25% variance. Timepoints follow a clockwise position, with distance between the first and last sampling.

Rhythmic detection was performed using MetaCycle (Table S3), and the identified rhythmic genes were categorized into different groups, as illustrated in Figure 2a. The total number of genes in each group is summarized in Figure 2b, while the specific genes in each group are detailed in Table S4. Out of Svevos’ 66,559 genes, we filtered out those with low expression (having fewer than 10 counts across all samples), leaving us with 46,975 genes. Among these, we identified 16,368 rhythmic genes under control conditions and 15,889 rhythmic genes under aphid infestation. Of these, 10,686 genes, referred to as the ‘Shared’ group, were found to be rhythmic in both control and aphid-infested conditions. The ‘Unique Control’ rhythmic genes group, comprising 5,682 genes, consists of those exhibiting rhythmicity only under control conditions. This group consists of genes that lost their rhythmicity during aphid infestation. The ‘Unique Aphid’ group, composed of 5,203 genes, comprises those that exhibit rhythmicity solely in response to aphid presence, having gained their rhythmic pattern as a result of infestation. Interestingly, the number of genes that lost rhythmic activity (5,682) and those that gained rhythmic activity (5,203) during aphid infestation are nearly equal.

**Figure 2.**
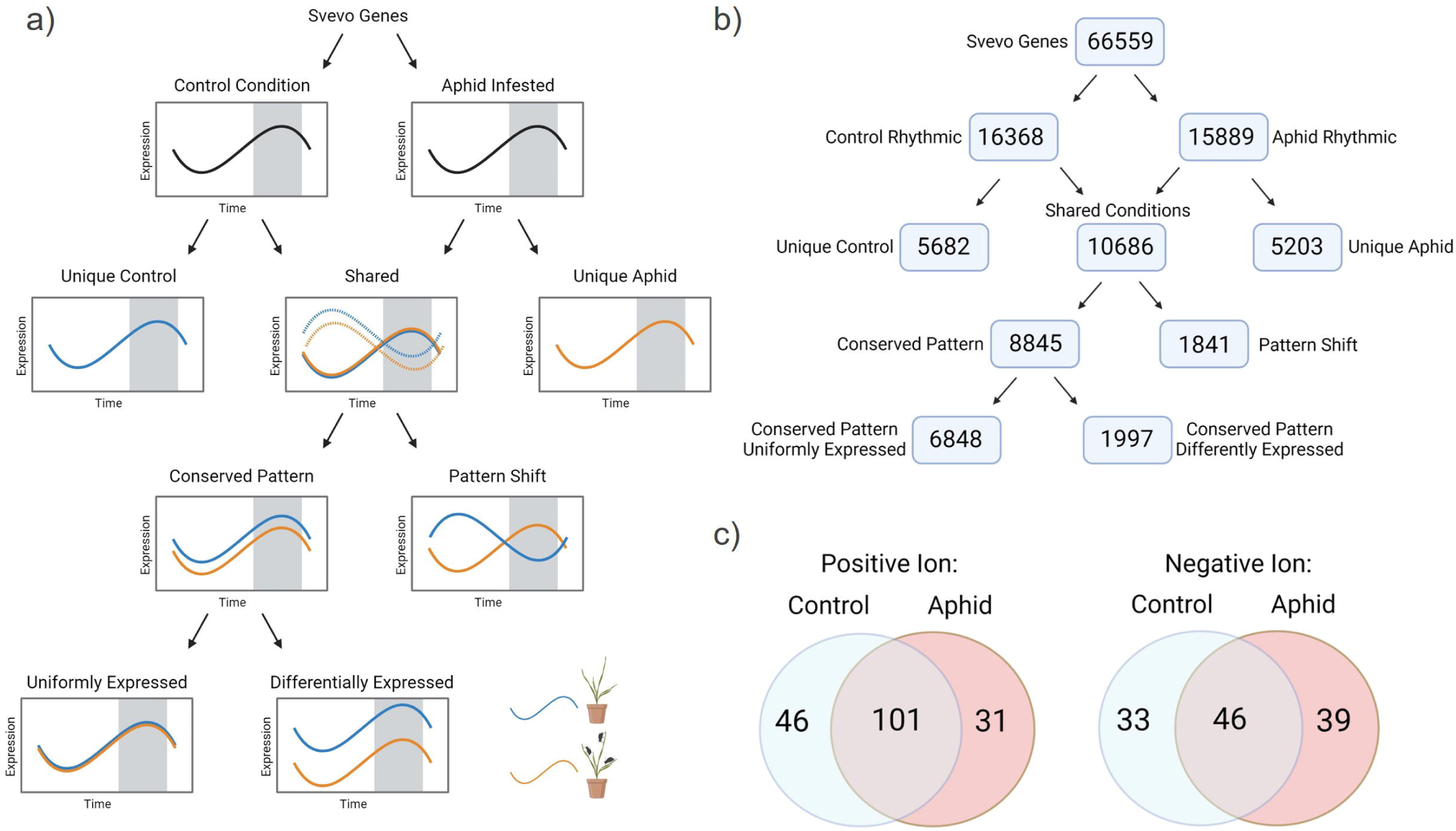
Classification of Svevo genes and metabolites into different rhythmic groups. (a) Division of Svevo genes into different rhythmic groups. Rhythmic detection was performed separately for control and aphid conditions using the Metacycle R package, with a corrected P-value cutoff of 0.05. Genes were grouped into those shared between both conditions and those unique to either control or aphid conditions, labeled as ‘Shared’, ‘Unique Control’, and ‘Unique Aphid’ respectively. Shared genes were further divided into those that maintained their rhythmic pattern and those that exhibited a shift, labeled as ‘Conserved Pattern’ and ‘Pattern Shift’ respectively. Finally, conserved pattern genes were split into differentially expressed under aphid infestation and uniformly expressed genes, labeled as ‘Conserved Pattern DE’ and ‘Conserved Pattern UE’ respectively. Differential expression analysis was conducted using DESeq2 with a corrected P-value cutoff of 0.05. (b) The number of rhythmic genes found in each of the different rhythmic groups. (c) Venn diagrams showing the number of rhythmic features detected in the control and aphid conditions, analyzed using untargeted metabolomics. Each diagram displays the number of rhythmic features in the ‘Unique Control’, ‘Shared’, and ‘Unique Aphid’ rhythmic groups, with positive and negative ion samples processed separately. Rhythmic detection was performed on the MS1 data obtained from LC-MS.

For genes found to be rhythmic in both conditions, we aimed to discern potential shifts in rhythmic expression patterns. The ‘Conserved Pattern’ group includes 8,845 genes that maintained their rhythmic pattern, while the ‘Pattern Shift’ group includes 1,841 genes that altered their rhythmic pattern.

We further divided the ‘Conserved Pattern’ rhythmic genes into differentially expressed (DE) and uniformly expressed (UE) groups. In total, 6,848 genes were identified as ‘Conserved Pattern UE’, while 1,997 genes were identified as ‘Conserved Pattern DE’. The ‘Conserved Pattern UE’ genes are those rhythmic genes whose pattern and expression level remain unaltered upon aphid infestation. Thus, approximately 40% of the rhythmic genes found under control conditions are minimally affected by aphid infestation. The ‘Conserved Pattern DE’ group is comprised of genes that under infestation exhibited a similar rhythmic pattern, but with either increased or decreased expression level.

### Rhythmic metabolite class composition alterations during aphid infestation

Rhythmic patterns were detected for both control and aphid-infested conditions using positive and negative ions modes separately with an LC-MS. The rhythmic metabolites were then categorized into the ‘Unique Control’, ‘Unique Aphid’, and ‘Shared’ groups, in a manner similar to the categorization performed for gene rhythmicity. For the ‘Unique Control’ and ‘Unique Aphid’ conditions, we identified 33 and 39 rhythmic features, respectively, under the negative ion mode, and 46 and 31 rhythmic features, respectively, under the positive ion mode (Figure 2c). The number of ‘Shared’ rhythmic features detected was 101 for the positive ion mode and 46 for the negative ion mode. The number of rhythmic features detected was determined from the acquired MS1 data, while the putative classification of rhythmic metabolites was determined from the MS2 data (Table S5).

The Natural Products Classification (NPC), obtained through the Sirius program, categorized the rhythmic metabolites into 10 unique superclasses (Figure 3). Superclasses, as defined by the NPClassifier program, represent broad categories of metabolites, including general classes of metabolites, chemical compounds, or biosynthetic information (Kim *et al*., 2021). The ‘Unique Control’ group contained a high percentage of small peptides (27%) and nucleosides (20%). Saccharides, found exclusively in the ‘Shared’ group, compose a high percentage (23%) of its rhythmic metabolomic composition. The ‘Unique Aphid’ group contains a high percentage of nucleosides (33%), aminosugars (13%), and monoterpenoids (10%), and a low percentage of small peptides, glycerolipids, and other metabolites (7%). A shift in the rhythmic metabolite composition is apparent under aphid infestation, as demonstrated by the varying compound classes.

**Figure 3.**
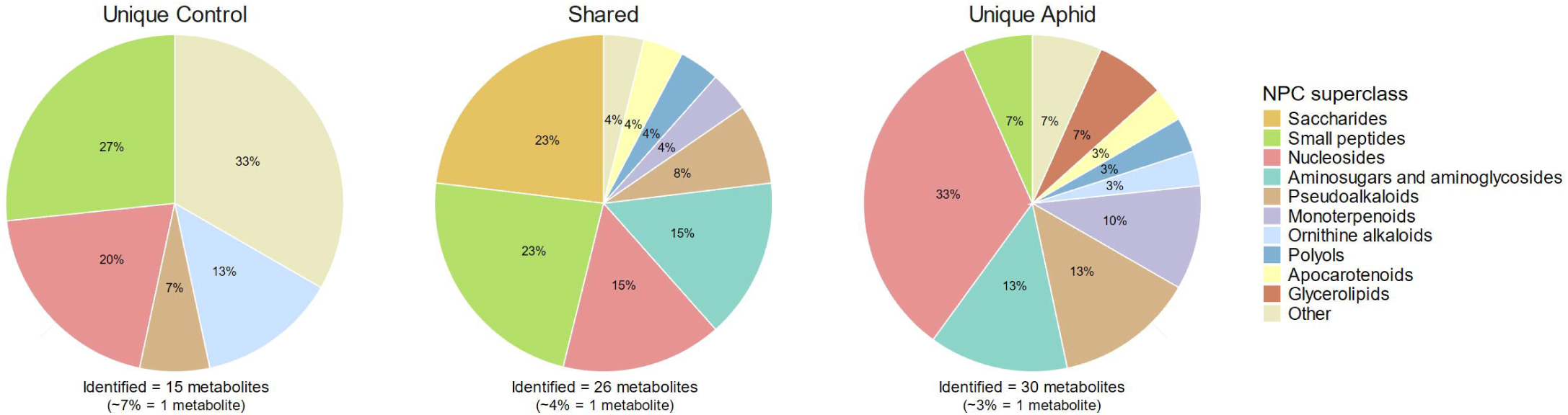
Rhythmic metabolites putative classification. Pie charts showing putative metabolite classification for the ‘Unique Control’, ‘Shared’, and ‘Unique Aphid’ rhythmic groups. Identification was performed on the MS2 data obtained from LC-MS using the CANOPUS tool within Sirius. Metabolites are classified by their Natural Product Compound (NPC) superclass family, as determined by NPClassifer.

### Gene Ontology (GO) enrichment reveals altered functions and processes under aphid infestation

GO enrichment was performed to observe distinct molecular functions and biological processes for the categorized rhythmic groups (Table S6). The ‘Unique Control’ group was enriched for vesicle-mediated transport and several binding functions, including GTP, RNA, fatty-acyl-CoA, and clathrin light chain binding (Figure 4a). The ‘Unique Aphid’ genes were enriched for defense responses, including response to abscisic acid (ABA) and heat shock protein 70 (Hsp70) protein binding (Figure 4b). The ‘Conserved Pattern’ group was enriched for photosynthesis and circadian rhythms (Figure 4c), as mentioned previously this group is further divided into differently expressed and uniformly expressed genes. The ‘Pattern Shift’ group, genes with a shifted rhythmic pattern, was enriched for phosphate ion transport and protein tyrosine kinase activity (Figure 4d). The circadian rhythm GO term was enriched in the ‘Conserved Pattern UE’ group (Figure 4e). Transmembrane transporter activity was enriched in both the ‘Conserved Pattern UE’ and the ‘Conserved Pattern DE’ rhythmic groups (Figure 4e, 4f), suggesting an important diurnal role for these transporters, whose rhythmic pattern is maintained under aphid infestation. The ‘Conserved Pattern DE’ group was highly enriched for down-regulated genes associated with photosynthesis activity, indicating a reduction in the gene expression, but not in the rhythm, of photosynthesis-related genes. The ‘Conserved Pattern DE’ group was also enriched for several metabolic processes, including carbohydrate and chlorophyll metabolic processes, and for cysteine and cellulose biosynthetic processes (Figure 4f).

**Figure 4.**
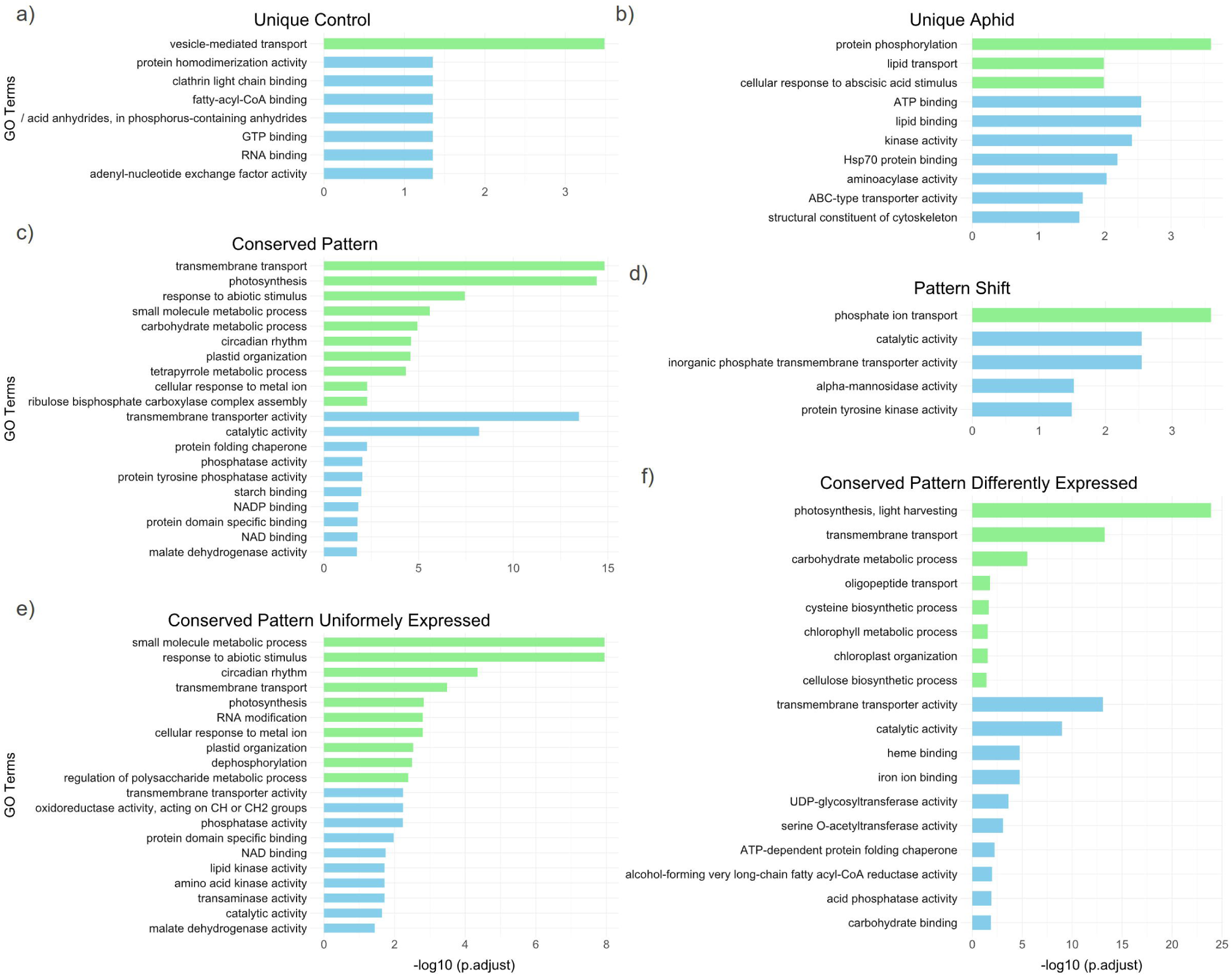
Gene Ontology (GO) term enrichment of different rhythmic gene groups. GO term enrichment of the different rhythmic groups was performed using the genes found in each group. The analysis was conducted on gProfiler using the Triticum turgidum genome from PlantEnsembl as the reference, with a BH-FDR cutoff of 0.05. Shown are the top 10 most significant GO terms for Biological Processes (green) and Molecular Functions (blue). The different rhythmic groups and their GO terms are: (a) Unique Control, (b) Unique Aphid, (c) Conserved Pattern, (d) Pattern Shift, (e) Conserved Pattern Uniformly Expressed, and (f) Conserved Pattern Differentially Expressed rhythmic groups. The symbol “/” stands for “hydrolase activity, acting on.

### Enriched transcription factor (TF) families in the categorized rhythmic groups

As regulatory proteins, TFs play crucial roles in the plant’s physiological functions. Identifying over or underrepresented transcription factor families can offer insights into the plant’s physiological adaptations. Therefore, we performed a TF family enrichment for the different rhythmic groups. TFs were acquired by running the plantTFDB transcription factor prediction tool (Table S7). TF enrichment was detected only for the ‘Shared,’ ‘Conserved Pattern,’ ‘Conserved Pattern DE,’ and ‘Conserved Pattern UE’ rhythmic groups.

The ‘Shared’ and ‘Conserved Pattern’ groups were both enriched for several TF families including CO-like, WRKY and Double B-box zinc finger (DBB), while only the ‘Shared’ group was enriched for HSF, auxin response factor (ARF), and B3 TF families (Figure 5 and Table S7). In the ‘Conserved pattern DE’ group, we observed an overrepresentation of rhythmic CO-like TF and the stress related HSF TF families. In the ‘Conserved pattern UE’ group, the DBB TF family was overrepresented, while the WRKY TF family was underrepresented. These findings highlight the dynamic regulatory roles of TF families in maintaining rhythmic patterns and responding to aphid-induced stress in plants.

**Figure 5.**
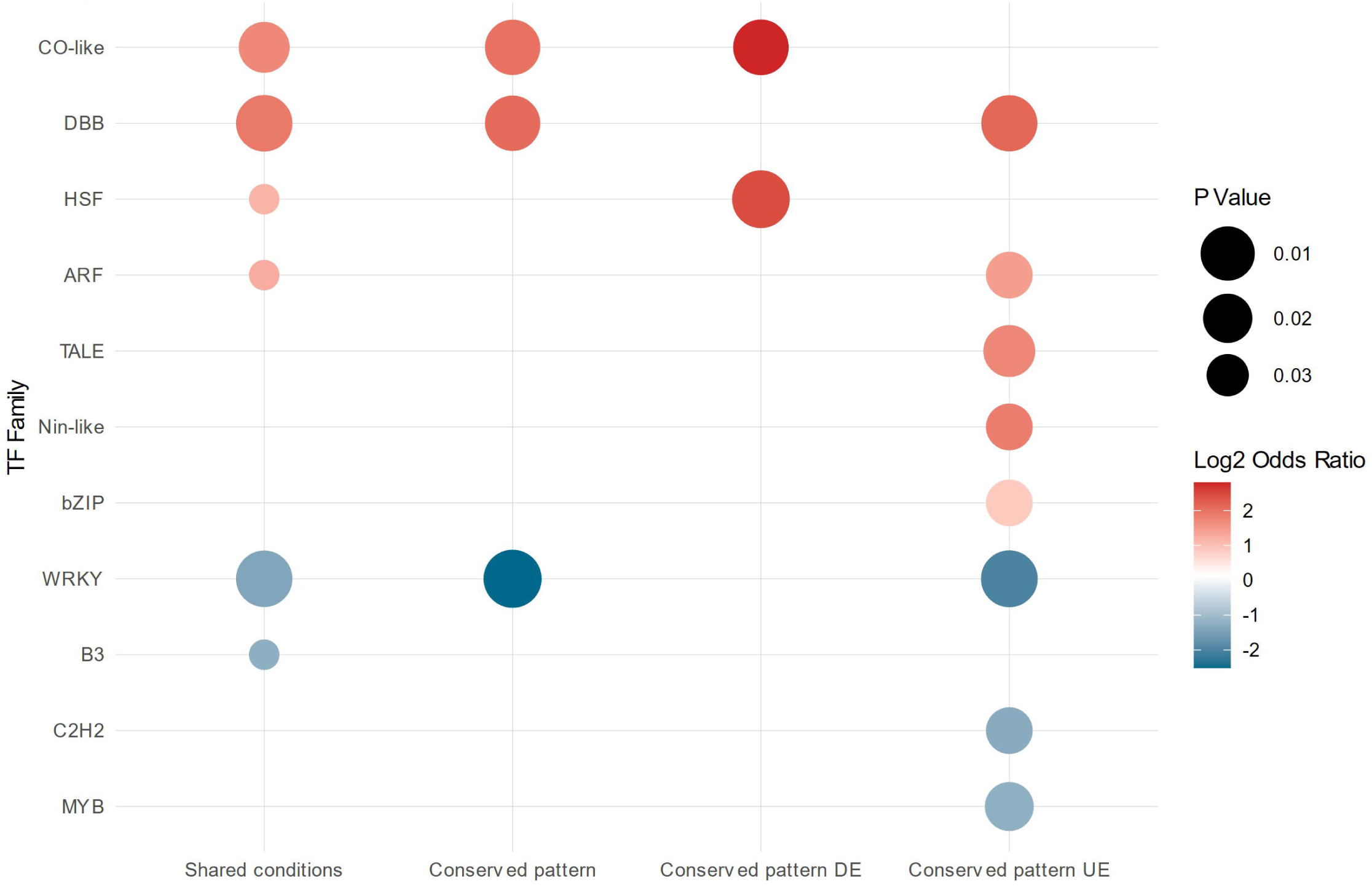
Transcription factor family enrichment for the different rhythmic groups. Transcription factor (TF) enrichment analysis was performed for all identified rhythmic groups. Gene TFs were first identified using the PlantTFDB TF prediction tool. Then, for each identified TF family, enrichment was tested for each rhythmic group, using all identified TF as background. Enrichment was observed only for the ‘Shared’, ‘Conserved Pattern’, ‘Conserved Pattern DE’ and ‘Conserved Pattern UE’ groups. Log2odds ratios are provided to show overrepresented and underrepresented TF families. A Fisher exact followed by Benjamin-Hochberg (BH) correction with a 0.05 cutoff was used to determine enrichment.

### Categorized rhythmic groups show variation in motif enrichment

Motif enrichment can provide details about which transcription factors may be regulating gene expression. By detecting enriched motifs, we can identify enriched binding sites for regulatory transcription factor in our rhythmic genes. Motif enrichment was performed on all the categorized rhythmic groups (Table S8). The top ten enriched motifs of the ‘Unique Control’, ‘Shared’, and ‘Unique Aphid’ rhythmic groups are presented in Table 1.

**Table 1.** Motif enrichment of different rhythmic groups. Motif enrichment was performed for all identified rhythmic groups using the AME tool from the MEME suite on the promoter (1 kb) sites of rhythmic genes. The AME tool was run using the JASPAR DNA Core Plants database as a motif reference. Here, the top 10 enriched motifs (with the most significant P values) are shown for the ‘Shared’, ‘Unique Control’, and ‘Unique Aphid’ rhythmic groups. The P values were calculated using the Fisher exact test followed by a Bonferroni correction with a 0.05 cutoff.

| **Condition** | **ID** | **Alt ID** | **p-value** |
| --- | --- | --- | --- |
| **Shared** | MA0972.1 | CCA1 | 1.06E-22 |
|  | MA1185.1 | LHY | 2.74E-19 |
|  | MA1401.1 | RVE7 | 1.65E-18 |
|  | MA1187.1 | RVE4 | 2.84E-17 |
|  | MA1183.1 | RVE6 | 1.18E-15 |
|  | MA1182.1 | RVE8 | 6.51E-15 |
|  | MA1834.1 | Zm00001d034298 | 1.95E-14 |
|  | MA1190.1 | RVE5 | 2.57E-14 |
|  | MA1184.1 | RVE1 | 2.24E-12 |
|  | MA1191.1 | RVE7L | 1.52E-10 |
| **Unique Control** | MA1753.1 | ERF057 | 4.67E-05 |
|  | MA1063.1 | TCP19 | 0.000094 |
|  | MA1240.1 | ERF10 | 0.000148 |
|  | MA1236.1 | ERF003 | 0.000169 |
|  | MA1833.1 | Zm00001d049364 | 0.000203 |
|  | MA1239.1 | ERF104 | 0.000229 |
|  | MA1062.2 | TCP15 | 0.000286 |
|  | MA1050.1 | OsI_08196 | 0.00034 |
|  | MA1019.1 | Glyma19g26560.1 | 0.000389 |
|  | MA1066.1 | TCP23 | 0.000723 |
| **Unique Aphid** | MA1805.1 | WRKY53 | 8.53E-08 |
|  | MA1088.1 | WRKY48 | 6.62E-07 |
|  | MA1078.1 | WRKY2 | 0.000026 |
|  | MA1084.1 | WRKY38 | 2.85E-05 |
|  | MA1077.1 | WRKY18 | 4.53E-05 |
|  | MA1086.1 | WRKY43 | 6.53E-05 |
|  | MA1075.1 | WRKY12 | 0.000147 |
|  | MA1090.1 | WRKY60 | 0.000736 |
|  | MA1093.1 | WRKY75 | 0.000934 |
|  | MA0589.1 | WRKY1 | 0.0026 |

The ‘Unique Control’ rhythmic genes were enriched for motifs from the ethylene response factor (ERF), TEOSINTE BRANCHED 1, CYCLOIDEA, and PCF1 (TCP) families. The ‘Shared’, ‘Conserved Pattern UE’, and the ‘Conserved Pattern DE’ groups were all enriched with motifs associated with the plants’ circadian rhythms. This includes the Circadian Clock Associated 1 (CCA1), REVEILLE (RVE), and LATE ELONGATED HYPOCOTYL (LHY) TFs (Table S8). The enrichment of circadian motifs in both the ‘Conserved Pattern UE’ and the ‘Conserved Pattern DE’ groups suggests that the core machinery controlling the timing of these rhythmic genes remains intact during aphid infestation. However, the presence of these motifs within the ‘Conserved Pattern DE’ indicates a potential regulatory role for the circadian clock in defense against aphid attacks.

In the ‘Unique Aphid’ group, only motifs associated with the WRKY TF family were enriched. WRKY TFs are known to play an important role in both biotic and abiotic responses. For instance, the Arabidopsis WRKY53 TF has been found to regulate plant defenses through jasmonic acid regulation (Jiao et al., 2022).

### Identifying rhythmic patterns of known circadian genes

Circadian genes are the molecular regulators of the plant internal clock; therefore, we expected them to show rhythmicity in our diurnal experiment. Circadian genes in Svevo were identified through homology to previously published circadian genes in *Triticum aestivum* (Rees et al., 2022). Most of the identified circadian genes were found in the ‘Conserved Pattern UE’ group, revealing that they maintained their expression under aphid infestation (Table. 2). Several circadian genes are found in the ‘Conserved Pattern DE’ group, including LUX_1B and RVE2/7 genes, suggesting that some circadian genes expression level are altered under aphid infestation. Additionally, two circadian genes ELF3 and CHE were found to be rhythmic only during aphid infestation.

### Rhythmic gene clustering and identification of hub transcription factors with their corresponding TF-gene-metabolite correlation network

Rhythmic genes from both aphid and control conditions were separated into clusters using Weighted Gene Co-expression Network Analysis (WGCNA). The rhythmic genes in control plants were grouped into eight clusters, while the rhythmic genes in the aphid-infested plants were grouped into 11 clusters. The expression of six clusters containing the highest number of genes is presented in Figure 6. The entire cluster list is detailed in Table S9.

**Figure 6.**
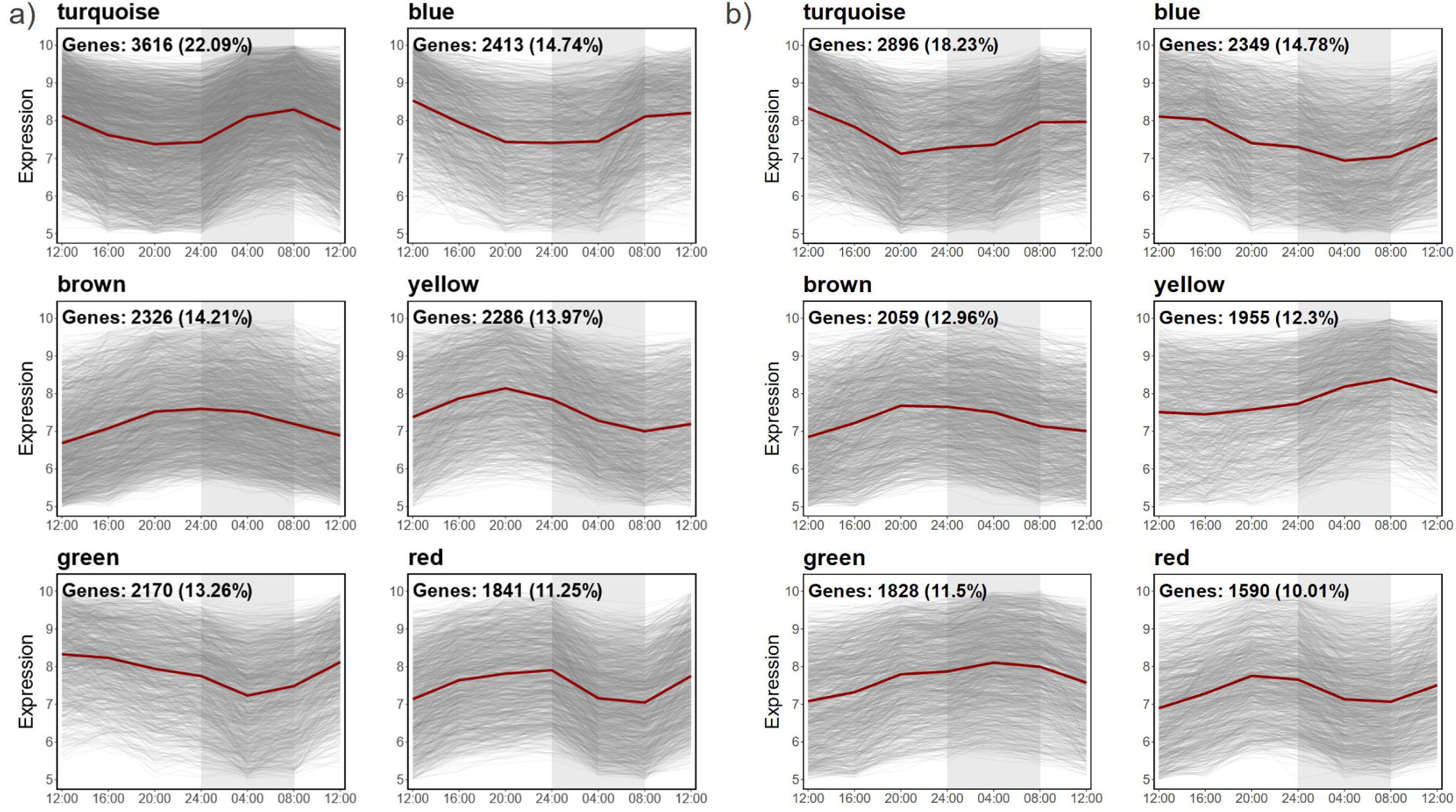
Rhythmic patterns of WGCNA clusters under control and aphid conditions. Rhythmic genes were clustered using WGCNA, applied separately to aphid and control conditions. Eight clusters were identified for the control condition and eleven clusters for the aphid condition. Shown here is the gene expression level of the top six clusters (those with the most abundant number of genes) for the (a) control and (b) aphid-infestation conditions. Gene expression levels were normalized with DESeq2 and then VST transformed prior to clustering. Each grey line represents a single gene and red lines indicate the cluster’s average. All gene clusters with corresponding genes can be found in Table S8.

To pinpoint activated pathways, we selected hub TF genes, defined as the TFs with the highest module membership within their respective cluster. All WGCNA clusters identified were used for hub TF identification. This analysis was carried out for both the ‘Unique Control’ and ‘Unique Aphid’ groups, identifying 14 hub TFs for the ‘Unique Control’ group and 19 for the ‘Unique Aphid’ group (Table S10).

We then identified correlation patterns between these hub TFs and rhythmic genes through a co-expression network analysis. This analysis, performed using CoExpNetViz, involved creating networks between the identified hub TFs and rhythmic genes. The co-expression networks where GO terms were detected are shown in Figures 7a and 7b, with the specific GO enrichment terms detailed in Table S10.

**Figure 7.**
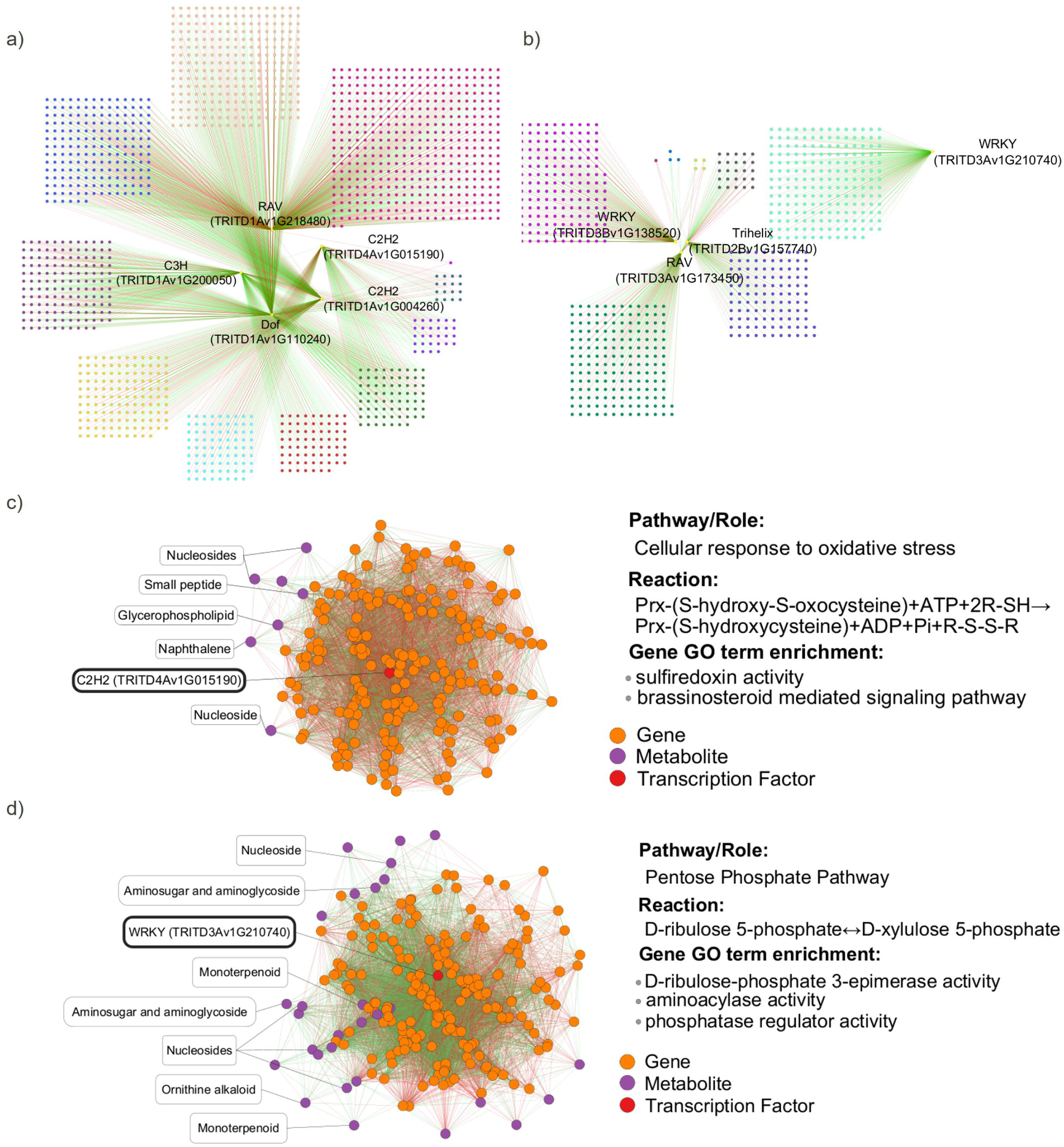
TF-gene and TF-gene-metabolite correlation networks. (a)-(b) Hub TF-gene correlation networks were created for the ‘Unique Control’ and ‘Unique Aphid’ groups. First, transcription factors belonging to the ‘Unique Control’ and ‘Unique Aphid’ groups were identified. Among these, hub TF genes were selected based on having the highest module membership in the different clusters (Table S8) identified using WGCNA. PCC co-expression networks were then created using the CoExpNetViz Cytoscape plug-in, correlating these hub TFs and rhythmic genes. Only networks with corresponding GO term enrichment (Table S9) for the identified correlated genes are shown. Nodes represent correlated genes and edges represent correlations, with green indicating a positive correlation and red indicating a negative correlation. Genes are grouped into their PLAZA family. (a) Hub TF and gene correlation network for the ‘Unique Control’ rhythmic group. (b) Hub TF and gene correlation network for the ‘Unique Aphid’ rhythmic group. (c)-(d) Hub TF-gene-metabolite PCC networks were created. The TFs and genes identified from the hub TF-gene CoExpNetViz correlation network were selected. These, together with the rhythmic metabolites in the ‘Unique Aphid’ and ‘Unique Control’ conditions, were used to create the networks. PCC networks were created using the ExpressionCorrelation Cytoscape plug-in. GO enrichment of the correlated genes was performed in gProfiler. Green edges indicate a positive correlation while red edges indicate a negative correlation. Two networks were created, one for (c) the C2H2 (TRITD4Av1G015190) of the ‘Unique Control’ group and the other for (d) the WRKY (TRITD3Av1G210740) of the ‘Unique Aphid’ rhythmic group.

A transcriptomics-metabolomics network can provide valuable insights into the functional relationships between genes and metabolites. Therefore, we created a correlation network from these hub TFs, their co-expressed rhythmic genes, and correlated rhythmic metabolites using ExpressionCorrelation (Figure 7c and 7d). In the correlation network analysis of the ‘Unique Control’ group, genes correlated with TRITD4Av1G015190, a C2H2 transcription factor, were linked to sulfiredoxin activity (Figure 7c). Sulfiredoxins are an important component of the plants redox system, contributing to the reduction of oxidative stress through the restoration of over-oxidized antioxidants. The network also included three nucleosides, a small peptide, a naphthalene, and a glycerophospholipid, all of which correlated with these genes.

In the ‘Unique Aphid’ group, the correlation network of TRID3Av1G210740, a WRKY transcription factor, showed enrichment associated with the pentose phosphate pathway (Figure 7d). The pentose phosphate pathway (PPP) is a major metabolic pathway involved in the production of NADPH and is one of the most important antioxidant cellular response systems. In addition, the PPP produces sugars that serve as precursors for nucleotide biosynthesis and aromatic amino acids. The correlated network contained four nucleosides, two amino sugars, two monoterpenoids, and an ornithine alkaloid.

### Transcriptome and metabolome integration for the ‘Unique Aphid’ rhythmic group

To further explore transcriptome and metabolome integration, the sPLS method from the mixOmics package was utilized. This method optimizes gene and metabolite groups by maximizing covariance and establishing correlations between them. A network integrating genes and metabolites for the ‘Unique Aphid’ group with a 0.7 correlation cutoff is shown in Figure S11.

The sPLS correlation between genes and metabolites identified two separate gene-metabolite hubs. GO term enrichment revealed that the identified genes were enriched in categories related to plant regulation and signaling, including protein phosphorylation and phospholipid transport (Xue et al., 2009; W., J., Zhang et al., 2023). Additionally, enrichment was observed for 3-hydroxyisobutyryl-CoA hydrolase activity, an enzyme linked to the valine catabolism pathway (Table S6 and Figure S11). The metabolites were classified into their respective NPC Superclasses. Among the correlated metabolites identified were a monoterpenoid and two pseudoalkaloids. Alkaloids and monoterpenoids, both classes of secondary metabolites, are known to contribute to plant defenses (Nagegowda et al., 2010; Koch et al., 2020).

## Discussion

### Under aphid infestation, many genes either gain or lose diurnal rhythmicity

Plants must constantly regulate their genes to optimize growth and adapt to changes between day and night. To elucidate the effect of biotic stress caused by aphid infestation on the plant’s diurnal rhythmicity, we conducted a time-series transcriptomics and metabolomics study comparing infested plants to non-infested plants. In this study, under control conditions, approximately 25% of Svevo genes exhibited rhythmic activity. While under aphid-infested conditions, approximately 24% of the total genes showed rhythmic activity (Figure 2b). Interestingly, only 16% of Svevo genes maintained rhythmic activity across both conditions, indicating that a significant number of genes are uniquely rhythmic in one condition or the other.

One factor that possibly determines the diurnal rhythmicity of a plant’s transcripts and metabolites are herbivore-associated molecular patterns (HAMPs). Plants recognize their attacker by specific HAMPs, while the attackers deliver an array of effectors (Snoeck et al., 2022). Aphid salivary effectors are crucial for aphid fitness on host plants and the modulation of plant defense responses to aphids, such as changes in redox status, degradation of cell walls, and regulation of the defensive signaling pathways (Bos et al., 2010; Mondal, 2020).

A well-characterized aphid effector is salivary protein ApC002 of pea aphids (*Acyrthosiphon pisum*), which is essential for continuous phloem feeding (Mutti et al., 2008). Our recent study on the interaction between *R. padi* and wheat showed that RpC002 and many other putative salivary effectors exhibited rhythmic diurnal patterns (Han et al., 2024). Since the plant is highly responsive to effectors, and some aphid salivary effectors exhibit rhythmic patterns, it is plausible that these rhythmic effectors could trigger rhythmic gene patterns in the infested plant.

We identified 5,203 (8%) genes that gained rhythmic activity in response to aphid infestation. Whether this is primarily due to the plant’s need to maintain a rhythm under stress, a response to rhythmic aphid-effectors, or other diurnal cycles in the aphid is yet to be determined. Possibly, all three factors contribute to the high number of induced rhythmic genes under infestation; however, this requires further investigation.

Many genes that retain their rhythmicity in both control and aphid-infested conditions are altered by the presence of aphids. This consists of a pattern shift where genes retain rhythmic activity yet exhibit altered rhythmic patterns. This change may reflect a response to the unique diurnal patterns of the aphids. Meanwhile, genes that maintain the same rhythmic pattern under both conditions may exhibit different expression levels under aphid infestation compared to control conditions. The preservation of their pattern may be linked to diurnal environmental conditions, such as light and temperature, which are unaffected by the presence of aphids.

### Untargeted metabolomics indicates changes in the diurnal rhythmicity of metabolites

Rhythmic metabolomics revealed differences in class abundance among the ‘Unique Control’, ‘Shared’, and ‘Unique Aphid’ groups (Figure 3). The ‘Unique Aphid’ group, which gained rhythmicity under infestation, exhibits a high percentage of monoterpenoids. The protective role of monoterpenoids may explain why the plant increases their accumulation and induces their rhythmic production under attack (Heiling et al., 2010; Pichersky and Raguso, 2018). The ‘Shared’ group is unique in containing rhythmic saccharides. Saccharides, which play important roles in plant defense against aphids, plant signaling, and the circadian clock, retained their rhythmicity under aphid infestation (Hofmann et al., 2010; Singh et al., 2011; Bolouri Moghaddam and Van Den Ende, 2013). These findings highlight a compositional shift in rhythmic metabolites in response to aphid infestation, further demonstrating the plant’s adaptive strategies.

### GO Enrichment analysis indicates a rhythmic defensive response to aphids

Rhythmic genes in the ‘Unique Control’ group, which lost their rhythmicity after aphid infestation, show enrichment in signaling-related GO terms such as GTP binding. This suggests that under infestation, many signaling genes may lose their rhythmic activity to accommodate different physiological processes. Similarly, the ‘Unique Aphid’ group, composed of genes that gained rhythmicity, is enriched with signaling terms, including protein phosphorylation and kinase activity. This hints at a mechanism of rhythmicity induction under aphid infestation, possibly through the activation of genes via post-transcriptional phosphorylation.

In addition, the ‘Unique Aphid’ group contains many stress-related GO terms, such as heat shock proteins (Hsp), protein binding and cellular response to abscisic acid (ABA). Hsp, primarily known for their induction under heat stress, can also be triggered by biotic stresses, including aphids (Sun et al., 2018; Shrestha et al., 2024). The observed rhythmic activity in genes responding to ABA may provide further insights into the established link between high ABA levels and increased aphid tolerance (Chapman et al., 2018). This demonstrates that certain components of the plant’s defense mechanism, including responses to Hsp and ABA, develop rhythmic patterns when exposed to aphid infestation.

The ‘Conserved Pattern DE’ group (Figure 4f) is enriched for the photosynthesis, light harvesting GO term. While this suggests that the light/dark cycle remains undisturbed by aphid infestation, infestation leads to a reduction in the expression of photosynthesis-related genes, which aligns with previous reports indicating that aphids hinder photosynthesis activity (Nietupski et al., 2022). The ‘Conserved pattern UE’ is enriched with genes related to circadian rhythms, further supporting the notion that plant’s internal clock remains unaffected.

Both the ‘Conserved Pattern DE’ and the ‘Pattern Shift’ groups are enriched with the transmembrane transport GO term. Transmembrane transport plays a crucial role in the shift from plant growth to defense, as demonstrated by ABCG36, an ABC transporter that modulates the balance between these two processes (Aryal et al., 2023).

### TF family enrichment indicates conserved circadian regulation under infestation

Transcription factor family enrichment can shed light on the plant’s response mechanisms during aphid infestation (Figure 5). DBB TFs were enriched in the ‘Conserved Pattern UE’ group. Known to be governed by the circadian rhythm and reactive to light, DBB transcription factors naturally appear among genes with a conserved pattern (Kumagai et al., 2008).

The ‘Conserved Pattern DE’ enrichment for the CO-like and HSF TFs suggests specific regulatory responses under aphid infestation. CO-like TFs have been found to mediate the interaction between the plant’s circadian rhythm and growth processes. CO-like genes also play a central role in photoperiod response and light-mediated reactions (Wong et al., 2014; Ruta et al., 2020). Their overrepresentation suggests that although aphid infestation does not disrupt the plant’s internal clock, it may influence plant growth regulated by these rhythmic TFs genes.

While HSFs are primarily known for their role in plant abiotic stress reactions, some have been found to interact with the circadian clock, while others respond to aphid infestation (Kolmos et al., 2014; Xia et al., 2014; Guo et al., 2016).Their overrepresentation in the ‘Conserved pattern DE’ group indicates that although they retain their rhythmic patterns, expression levels of these stress related TF family are altered when plants are exposed to aphids, possibly in response to stress induced by the infestation.

### Motif enrichment revealed a WRKY specific response under aphid infestation

Motif enrichment was performed to identify potential regulators of rhythmic genes. Upstream regulatory motifs act as binding sites for transcription factors, which in turn regulate gene expression. Motif enrichment analysis for the ‘Unique Control’ group revealed several motifs, primarily from the ERF and TCP families (Table 1). ERF TFs are ethylene-sensitive transcription factors involved in plant growth as well as abiotic and biotic stress (Cheng et al., 2013; Wu et al., 2020; L., Zhang et al., 2020). ERF TFs can serve as intermediaries between jasmonic acid (JA) and ethylene (ET) signaling (Pré et al., 2008). Additionally, several ERF transcription factors depend on ABA for their function (Li et al., 2011). Given their roles as intermediaries in both JA-ET and ABA signaling pathways, ERF transcription factors are sensitive to hormone signaling and help initiate the plant’s defense mechanisms against stress (Cheng et al., 2013). TCP TFs, like ERFs, are also mediated by a range of hormones, including ABA, auxin, brassinosteroid, cytokinin, gibberellin, JA, and SA (Danisman, 2016). The TCP TF family is mostly known for its involvement in plant development (Danisman, 2016; Koyama et al., 2017). However, TCP TFs can also enhance plant defenses. TCP 8/14/15/20/22/23, which are enriched in the ‘Unique Control’ group (Table S8), have been shown to interact with the effector-triggered immunity (ETI) negative regulator SRFR1 (Kim et al., 2014). Furthermore, several TCP TFs have been found to bind to the plants core circadian clock components, subsequently influencing their transcript levels (Giraud et al., 2010). Here, ERF and TCP TF families, which maintain rhythmic regulation under control conditions, lose their rhythmicity when reacting to aphid-induced stress. Given their dual roles in development and defense, their loss of rhythmic activity under infestation indicates a disruption in rhythmic processes related to both growth and defensive.

The ‘Shared’ group genes were enriched for motifs related to the plants circadian clock including CCA1, LHY, and RVE genes (Table 1). Circadian Clock Associated 1 (CCA1), a known central circadian clock regulator, also enhances defenses under aphid infestation (Lei et al., 2019). The ‘Conserved Pattern DE’ group was also enriched with several circadian genes, including CCA1 and RVE 4/6/8, reinforcing the notion that circadian genes are vital to plant defenses.

The ‘Unique Aphid group’, consisting of gained rhythmic genes under infestation, are enriched for WRKY motifs (Table 1). The WRKY transcription factor family is recognized for its involvement in plant defenses, with numerous WRKY genes observed to be induced during aphid infestation (Y., Zhang et al., 2020). WRKY transcription factor families contribute to plant protection through various methods, including callose formation, the production of secondary metabolites, plant signaling, and the modulation of hormone responses such as SA and JA (Schluttenhofer and Yuan, 2015; Kloth et al., 2016; Poosapati et al., 2022). As noted earlier, the ‘Unique Aphid’ group is enriched for the kinase activity GO term; one function of kinase activities is to regulate WRKY responses (Pandey and Somssich, 2009). It is worth noting that the WRKY TF motifs are exclusively enriched in this group, alluding to some role WRKY genes might play in relation to the plant rhythmic response to aphid infestation. Recently, connections between the circadian clock and WRKY transcription factors, as well as their regulation of downstream responses, have been unveiled. Wang *et al*. demonstrated that Arabidopsis WRKY2/10 TFs bind to CCA1/LHY and to Phytochrome Interacting Factor 4 (PIF4), a central growth regulator. This interaction, in turn, regulates the expression of key defensive genes (Wang et al., 2022). Boyte *et al*. identified multiple WRKY TFs involved in ROS-regulated sugar signaling, highlighting their connection to sugar-activated superoxide’s production and their effects on the circadian clock (Boyte et al., 2023). We speculate that under infestation, WRKY TFs bind to both core circadian clock and defense-related genes, thereby initiating an induced rhythmic response. However, further investigation is needed to reveal their exact role.

### Most core circadian clock genes maintain diurnal rhythmicity in response to herbivore infestation

The plants diurnal cycle is controlled by both external cues, such as sunlight and temperature, and by the plants internal clock. Most of the core circadian genes reside in the ‘Conserved pattern UE’ group, which is minimally affected by aphids, suggesting that these genes are largely unaltered by aphid infestation (Table 2). Interestingly, both RVE 2/7 and LUXlb are the only circadian genes that retained their rhythmic pattern but exhibited different expression levels. Although the role of circadian genes in plant defense is well documented, the specific function of the differentially expressed circadian genes in response to aphid infestation is not yet understood (Zhang et al., 2019; Butt et al., 2020). Notably, the circadian genes ELF3 and CHE were found to exhibit rhythmic activity only under aphid infestation, further investigation is needed to clarify this finding

**Table 2.** Core circadian genes categorization into rhythmic groups. Core circadian genes were identified as orthologues of known *Triticum aestivum* (bread wheat) circadian genes. Once identified, the circadian genes were categorized based on the rhythmic group they were found in. All identified core circadian genes were found in either the ‘Conserved Pattern UE’, ‘Conserved Pattern DE’, or the ‘Unique Aphid’ rhythmic group.

| **Gene** | **Suggested name** | **Common name** | **Rhythmic group** |
| --- | --- | --- | --- |
| **TRITD3Av1G037220** | TdGI_3A | GI | Conserved Pattern UE |
| **TRITD3Bv1G046120** | TdGI_3B | GI | Conserved Pattern UE |
| **TRITD7Av1G151430** | TdLHY_7A | LHY | Conserved Pattern UE |
| **TRITD7Bv1G108620** | TdLHY_7B | LHY | Conserved Pattern UE |
| **TRITD4Av1G112820** | TdLNK1_4A | LNK1 | Conserved Pattern UE |
| **TRITD4Bv1G100760** | TdLNK1_4B | LNK1 | Conserved Pattern UE |
| **TRITD3Av1G109260** | TdLNK2_3A | LNK2 | Conserved Pattern UE |
| **TRITD3Bv1G108050** | TdLNK2_3B | LNK2 | Conserved Pattern UE |
| **TRITD3Av1G282500** | TdLUX/BOA_3A | LUX/BOA | Conserved Pattern UE |
| **TRITD1Av1G164060** | TdLUX-Lb_1A | LUX-Lb | Conserved Pattern UE |
| **TRITD6Av1G099490** | TdLWD_6A | LWD | Conserved Pattern UE |
| **TRITD5Av1G182370** | TdPRR59_5A | PRR5/PRR99 | Conserved Pattern UE |
| **TRITD4Bv1G124260** | TdPRR73_4B | PRR3/PRR7 | Conserved Pattern UE |
| **TRITD6Av1G165480** | TdRVE68_6A | RVE6/RVE8 | Conserved Pattern UE |
| **TRITD6Bv1G150850** | TdRVE68_6B | RVE6/RVE8 | Conserved Pattern UE |
| **TRITD7Av1G249660** | TdRVE86b_7A | RVE6/RVE8 | Conserved Pattern UE |
| **TRITD6Bv1G133070** | TdTOC1_6B | TOC1 | Conserved Pattern UE |
| **TRITD1Bv1G149820** | TdLUX-Lb_1B | LUX-Lb | Conserved Pattern DE |
| **TRITD2Av1G258530** | TdRVE27a_2A | RVE2/RVE7 | Conserved Pattern DE |
| **TRITD2Bv1G220960** | TdRVE27a_2B | RVE2/RVE7 | Conserved Pattern DE |
| **TRITD7Bv1G230030** | TdRVE27b_7B | RVE2/RVE7 | Conserved Pattern DE |
| **TRITD6Bv1G152230** | TdRVE27c_6B | RVE2/RVE7 | Conserved Pattern DE |
| **TRITD6Av1G150650** | TdCHEb_6A | CHE | Unique Aphid |
| **TRITD1Av1G230270** | TdELF3_1A | ELF3 | Unique Aphid |

### Transcriptome and metabolome integration reveals plants molecular responses to aphid infestation

The ‘Unique Control’ group contained a correlation network between the hub TFs, their co-expressed rhythmic genes, and correlated rhythmic metabolites that was associated with sulfiredoxin activity and the brassinosteroid mediated signaling pathway. Sulfiredoxins assist in reducing peroxiredoxins, which are antioxidants crucial for reducing reactive oxygen species. Consequently, they contribute to restoring plants exposed to elevated oxidative stress. Brassinosteroid, a plant hormone, plays various roles in plant growth and development. Under stress, it assumes a role in modulating the balance between plant growth and immunity (Ortiz-Morea et al., 2020). It has been shown to bolster defenses by promoting H2O2 production and regulating glucosinolate biosynthesis (Xia et al., 2011; Lee et al., 2018).

The correlation network between the hub TF and their correlated rhythmic genes and metabolites in the ‘Unique Aphid’ group was associated with the pentose phosphate pathway (PPP), as indicated by the enrichment of ribulose phosphate 3-epimerase activity. NADPH, an energy storage molecule generated in the PPP pathway, functions in various plants processes, including redox signaling and ROS detoxification (Tripathy and Oelmüller, 2012).

Oxidative species can function in cell signaling and plant protection from pests; however, in excess, they are toxic (Goggin and Fischer, 2022). Therefore, to maintain optimal oxidative levels, they must be constantly regulated. Notably, oxidative balance responses, presented in both the ‘Unique Control’ and ‘Unique Aphid’ groups, suggest that oxidative levels must be tightly regulated both under control conditions and during aphid infestation.

Using sPLS for metabolomics and transcriptomics integration, we observed that the rhythmic genes showing the highest covariance with rhythmic metabolites are associated with plant signaling, including protein phosphorylation and phospholipid transport. These results highlight a link between plant signaling and rhythmic metabolite dynamics. We observed that two alkaloids, a monoterpenoid, and three unidentified metabolites exhibited the strongest covariance to the ‘Unique Aphid’ rhythmic genes. Alkaloids, a large family of secondary metabolites, are often toxic to insects. High alkaloid content in plants is linked to increased aphid resistance (Adhikari et al., 2012). Terpenes are another class of secondary metabolites; during infestation, plants increase terpene synthesis to attract the natural parasitoids of pests (Yuan et al., 2008).

## Conclusion

To summarize, we identified induced defense-related rhythmic activity in wheat under aphid infestation, with WRKY TF genes playing a key role. Core circadian genes maintained their rhythmic activity, and oxidoreductase levels remain tightly controlled. Furthermore, shifts in the rhythmic patterns of gene expression and metabolomic composition were identified, highlighting the trade-off between plant growth and defense responses. To cope with aphid attacks, the plant likely downregulates and alters rhythmic genes essential for growth while activating and modifying the expression levels of rhythmic genes involved in stress response. Although the plant must defend itself against aphid infestation, it cannot ignore the changes throughout the diurnal cycle and must maintain a rhythmic pattern that complements both its own daily cycle and the aphids’ diurnal cycle.

Our findings shed light on the complex plant-insect interaction within the plant diurnal cycle. Understanding these daily interactions can potentially enhance pest management strategies. Chronoculture is a practice that exploits the daily biological clock by aligning farming activities with natural daily cycles (Steed et al., 2021). Utilizing time of day specific pest management applications can help farmers improve yield. By recognizing and utilizing optimal times, farmers may enhance field farming practices to more effectively target harmful pests.

## Material and Methods

### Plant growth conditions and aphid colony

Tetraploid wheat *Triticum turgidum* ssp. *Durum* (cultivar Svevo) plants were cultivated in a growth tent under a 16:8 L:D light regime. The temperature was maintained at 24±2°C during light hours and 20±2°C during dark hours. Wheat was planted in a 98-conical pot array (Ray Leach Cone-tainers™). The experiment commenced 14 days post-germination, when the wheat seedlings were between the second and third leaf stages (Zadoks 12 and 13). The bird cherry-oat aphids (*R. padi*), were reared and age-synchronized for 6-days on two-week-old plants of *T. aestivum* cv. Chinese Spring.

### Plant-aphid diurnal rhythm experimental setup

Approximately 40 adult wingless aphids were placed on the second leaf of each Svevo plant. The aphids were kept inside a cage composed of a Falcon^®^ tube (Fisher Scientific, Waltham, MA, USA), with the side closed off with a cotton ball. Control plants, without aphids, also had a Falcon tube cage attached to ensure any potential cage effects were consistent across all plants. All aphids were positioned on the plants at 4zt (12:00). Every four hours, leaf areas containing aphids were collected from the aphid-infested plants, and comparable areas were taken from the control plants. Tissues were taken at each collection timepoint, continuing until 4zt (12:00 of the next day). All collected samples were immediately flash-frozen in liquid nitrogen and stored at −80°C until further processing for RNA and metabolite isolations (see scheme in Figure 1a). A total of four biological replicates were included for RNA sequencing (RNA-seq) analysis, and six biological replicates were included for metabolomic analysis. RNA-seq collection was conducted at Colorado State University (CSU), while metabolomic collection was conducted at Ben Gurion University (BGU).

### RNA extraction, library construction, and RNA sequencing

Plant RNA was extracted using the Norgen Total RNA extraction kit and treated with DNAse (Norgen Biotek, Thorold, Ontario, Canada), following the manufacturer’s protocol. RNA concentration was measured using a Nanodrop Spectrophotometer, and its integrity was confirmed with a Bioanalyzer prior to sequencing. Library construction and RNA sequencing was conducted at Novogene Corporation Inc. (Sacramento, California). The poly(A) mRNA-enriched libraries were constructed using NEBNext® Ultra™ II RNA Library Prep Kit (New England BioLabs, Ipswich, MA). Sequencing was performed using an Illumina NovaSeq 6000 platform, producing 150-bp paired-end reads with an average depth of 27.8 Mbp. Adapters were trimmed with TrimGalore v0.6.10 (Krueger, 2015) and then filtered to ensure quality reads. Reads were aligned to the Svevo.v1 reference genome (Maccaferri et al., 2019), sourced from Plant Ensemble, using STAR alignment(Dobin et al., 2013) with default settings generating gene counts (Table S1). RNA-seq quality was assessed by consolidating reports from FastQC v0.12.1 (Andrews, 2010) into a MultiQC v1.14 (Ewels et al., 2016) report and further analyzed using SeqMonk software (Andrew, 2007).

### RNA-seq data analysis

Genes with fewer than 10 reads across all timepoints were filtered out. Differential expression analysis was performed using the DESeq2 package v1.38 (Love et al., 2014) in *R* (Table S2). Criteria for differential expression were an absolute log_2_ fold change exceeding 1 and an adjusted Benjamin Hochberg *P*-value less than 0.05. Using the DESeq2 package, reads were normalized by estimating the size factor and then subjected to a Variance Stabilization Transformation (VST) for all subsequent analyses. A Principal Component Analysis (PCA) was generated using MetaboAnalyst v5.0 (Pang et al., 2021), applying a 25% Interquartile Range (IQR) filter and Auto-scaling. Gene Ontology (GO) term enrichment was conducted through gProfiler (Raudvere et al., 2019) via the g:COSt function. We used only the expressed annotated genes as a background, drawing from the GO terms for *Triticum Tugurium* in Plant Ensemble. An adjusted *P*-value cutoff of 0.05 was used for enrichment analysis and only highlighted terms were kept. Highlighted terms are those identified through gProfiler’s filtering algorithm, which retains only the most significant GO terms. Results from the enrichment analysis were visualized using the ggplot2 package (Wickham, 2011) in *R*.

### LC-MS metabolic profiling

Six replicates of plant tissue (leaves) were collected for metabolomic analysis and stored as stated above. The tissues were then grinded, and 50 mg of the fine powder was diluted in a solvent composed of 80% methanol and 0.1% formic acid, with 1,3-benzoxazol-2-one (BOA) at 10 µg/ml serving as an internal standard. Samples were diluted with the extraction solvent in a 1:10 (mg: µL) ratio.

Ultra-performance liquid chromatography coupled with high-resolution mass spectrometry (UPLC-HRMS) analysis was conducted at the Metabolomics unit of the Ilse Katz Institute for Nanoscale Science & Technology of BGU.

Samples were subjected to metabolite extraction, focusing on (semi)polar compounds (Batushansky et al., 2019), and were run through Waters Acquity LC equipped with an HSS T3 reverse-phase column, connected to Thermo Q-Exactive MS equipped with an electron spray ionization (ESI) source. The gradient of mobile phases A (0.1% formic acid in water) and B (0.1 formic acid in acetonitrile) was applied as follows: 0-1 min 99% of A, 1-11 min 60% of A, 11-13 min 30% of A, 13-15 min 1% of A, 15-16 min 1% of A. 16-17 min 99% of A, 17-20 min 99% of A. The flow rate was 0.4 mL/min, and the column temperature was maintained at 40°C. Data acquisition was performed in full scan (60-1000 m/z) positive and negatives modes. Additionally, to the samples, three quality control pools were analyzed in both full scan and MS/MS modes. Subsequently, raw files were processed using the MSConvert tool of ProteoWizard v3.0 (Kessner et al., 2008).

### Metabolites data analysis

The mass spectrometry data were first processed with MZMINE3 (Schmid et al., 2023) for peak detection, alignment, and quantification. The results were then exported to Global Natural Products Social Molecular Networking (GNPS) for Feature-Based Molecular Networking (FBMN) analysis (Wang et al., 2016; Nothias et al., 2020). The data was filtered by removing all MS/MS fragment ions within +/− 17 Da of the precursor m/z. MS/MS spectra were window filtered by choosing only the top 6 fragment ions in the +/− 50 Da window throughout the spectrum. The precursor ion mass tolerance was set to 0.05 Da and the MS/MS fragment ion tolerance to 0.05 Da. A molecular network was then created with the FBMN workflow on GNPS. The network was created where edges were filtered to have a cosine score above 0.70 and more than 6 matched peaks. Further, edges between two nodes were kept in the network if and only if each of the nodes appeared in each other’s respective top 10 most similar nodes. Finally, the maximum size of a molecular family was set to 100, and the lowest scoring edges were removed from molecular families until the molecular family size was below this threshold. The spectra in the network were then searched against GNPS spectral libraries (Horai et al., 2010; Wang et al., 2016). The library spectra were filtered in the same manner as the input data. All matches kept between network spectra and library spectra were required to have a score above 0.7 and at least six matched peaks. The DEREPLICATOR was used to annotate MS/MS spectra (Mohimani et al., 2018).

Compound classification was conducted using Sirius (Dührkop et al., 2019) and CSI:FingerID (Dührkop et al., 2015; Hoffmann et al., 2021). The mass spectrometry data were integrated into Sirius v 5.8.0 for identification. CSI:FingerID, within Sirius, predicted the structures of unidentified compounds by contrasting the obtained spectra with *in silico* fragmented spectra from public databases. Subsequently, CANOPUS (Dührkop et al., 2021), also within Sirius, facilitated the categorization of compounds into their respective NPClassifier (Kim et al., 2021) classes.

Features were then filtered against both blank samples and the coefficient of variation (CV) of quality control samples. After filtering, the features were log_2_ normalized prior to rhythmic detection.

### Rhythmic detection

Gene rhythmicity was assessed using the meta2d program of the MetaCycle (Wu et al., 2016) *R* package, this is a tool that employs several common statistical tests for detecting rhythms. In this study, both LS and JTK statistical rhythmic tests were conducted using MetaCycle, and their results were combined through the Bonferroni method. An adjusted Benjamin Hochberg *P*-value of 0.05 served as the cutoff to identify rhythmicity. All results were validated by testing for rhythmicity using Nitecap (Brooks et al., 2022).

Rhythmic detection was performed for both genes under control and aphid-infested conditions. Genes were divided into different groups: genes rhythmic in both conditions were placed in the ‘Shared’ group, genes rhythmic only under control conditions in the ‘Unique Control’ group, and genes rhythmic only under aphid-infested conditions in the ‘Unique Aphid’ group. To identify genes that maintain a rhythmic pattern, but with a potential pattern shift, the PCC was determined between its expression under control and aphid-infested conditions. Specifically, for each gene found to be rhythmic in both conditions, the PCC was determined between its expression under control and aphid-infested conditions. A 0.7 correlation cutoff was used to determine conserved patterns (‘Conserved Pattern’) and pattern shifts (‘Pattern Shift’). Additionally, to identify genes with a conserved pattern under infestation but with differential expression, Deseq2 was employed. All rhythmic genes that were differently expressed (DE) at least at one timepoint between control and aphid-infested conditions were considered DE. Using this approach, genes were categorized as ‘Conserved Pattern DE,’ or as ‘Conserved Pattern UE.’

Metabolite rhythmicity under control and aphid-infested conditions was similarly identified using Metacycle’s metad2d program and validated with Nitecap on the normalized feature data. The NPClassifier Superclass distribution of rhythmic metabolites was visualized in a pie chart via the ggplot2 package in *R*.

### Transcription factor identification and enrichment

Transcription factors (TFs) in Svevo were identified using PlantTFDB v5.0 (Jin et al., 2017), a server that predicts TFs by comparing protein sequences to established transcription factors. For every rhythmic group, including the ‘Unique Control’, ‘Unique Aphid’, ‘Shared’, ‘Conserved Pattern’, ‘Pattern Shift’, ‘Conserved Pattern UE’, and ‘Conserved Pattern DE’ groups TF enrichment was analyzed, using all expressed genes as the background. TF enrichment was determined using a Fisher Exact test, followed by a Benjamin-Hochberg (BH) correction with a cutoff of 0.05.

### Motif identification and enrichment

Using the AME program suite v5.5 (Bailey et al., 2009), motif enrichment of rhythmic genes across the different rhythmic groups, specifically the ‘Unique Control’, ‘Unique Aphid’, ‘Shared’, ‘Conserved Pattern’, ‘Pattern Shift’, ‘Conserved Pattern UE’, and ‘Conserved Pattern DE’ groups, was assessed. Promoter regions were defined as the 1 kb sequence located upstream of the coding regions of these rhythmic genes. The JASPAR DNA Core Plants database (2022) (Castro-Mondragon et al., 2022) served as the motif database. Motifs from genes exhibiting expression were utilized as the background reference.

### Identification of core circadian genes

Svevo orthologues of previously identified circadian *Triticum aestivum* genes (Rees et al., 2022), were acquired from Plant Ensemble (Bolser et al., 2016). Only genes with high confidence and high DNA similarity (>99.2%) were considered orthologues.

### Rhythmic gene clustering

Gene clustering was performed on VST-normalized expression levels using the Weighted Gene Co-Expression Network Analysis (WGCNA) package (Langfelder and Horvath, 2008) in *R*. Rhythmic genes from both aphid-infested and control conditions were analyzed separately using WGCNA. After establishing a scale-free topology, a soft power of 3 was determined for both the aphid and control conditions. The parameters used for WGCNA were: networkType = “signed”, minModuleSize = 30, corType = “bicor”, maxBlockSize = 15000, TOMType = “signed”, power = soft_power, and mergeCutHeight = 0.25. The gene clusters and their corresponding module memberships were then recorded.

### Transcription factor co-expression network analysis

Hub genes were pinpointed by selecting only the genes found in the top 0.05% highest module membership of the different clusters in the WGCNA analysis. These were further filtered to keep only TF genes, specifically those that were identified as TFs through PlantTFDB. A co-expression network was then created between the identified hub TFs and rhythmic genes of the ‘Unique Control’(those that lost rhythmic activity) and ‘Unique Aphid’(those that gained rhythmic activity) groups. This analysis was administered in Cytoscape v3.10.0 (Shannon et al., 2003) using the CoExpNetViz plugin (Tzfadia et al., 2016). CoExpNetViz establishes a network based on the Pearson Correlation Coefficient (PCC) between the “baits” (Hub TFs) and the “targets” (Rhythmic genes). The genes are categorized by gene families sourced from PLAZA (Van Bel et al., 2022) and are clustered using the Tribe-MCL Algorithm (Enright et al., 2002). In CoExpNetViz, strict cutoffs were set with the upper percentile at 99 for positive correlations and the lower percentile at 1 for negative correlations.

### Transcriptome and metabolome integration

The ExpressionCorrelation plugin in Cytoscape was used to integrate hub TFs, rhythmic genes, and rhythmic metabolites, applying a 0.7 correlation cutoff. The sPLS function of the mixOmics v6.24.0 *R* package was employed to integrate transcriptomics and metabolomics data (Rohart et al., 2017). Tuning was conducted to identify the optimal number of genes and metabolites for integration. Rhythmic genes were grouped in increments of 50 to 300, while metabolites were grouped in increments of 3 to 15. Cross-validation was performed by repeating the sPLS tuning process 100 times.

### Data availability

The raw MS metabolomic data has been deposited in the MassIVE database under accession number MSV000095493. The raw RNA sequencing data has been deposited in NCBI under accession number PRJNA1142710. The metabolites identifications and networks for the negative and positive ions using GNPS can be found on the GNPS website via: https://gnps.ucsd.edu/ProteoSAFe/status.jsp?task=7614750cd37b4f25abdaf2df4c9dc053 for the negative ion and https://gnps.ucsd.edu/ProteoSAFe/status.jsp?task=1266bb00035a4cbca1c5c4dc5edb1f9e for the positive ion.

## Acknowledgement

We would like to extend our sincere gratitude to Sariel Hubner (Galilee Research Institute, Tel-Hai College) and Beery Yaakov (Ben-Gurion University of the Negev) for their invaluable insights, constructive feedback, and thoughtful discussion.

